# Testing the auditory steady-state response (ASSR) to 40-Hz and 27-Hz click trains in children with autism spectrum disorder and their first-degree biological relatives: A high-density electroencephalographic (EEG) study

**DOI:** 10.1101/2025.08.05.668742

**Authors:** Megan Darrell, Theo Vanneau, Dennis Cregin, Tringa Lecaj, John J. Foxe, Sophie Molholm

## Abstract

**Motivation:** Altered auditory processing likely contributes to core social and attentional impairments in autism spectrum disorder (ASD). The auditory steady-state response (ASSR)— a neural measure of auditory processing and cortical excitatory-inhibitory balance—has yielded mixed results in ASD. This study uses high density electroencephalography (EEG) to evaluate ASSR in ASD and unaffected siblings to clarify neural mechanisms underlying auditory deficits in autism.

**Methods:** High-density 70-channel EEG was recorded in children (8–12 years, IQ >80) with ASD (n=53), typically developing (TD) peers (n=35), and unaffected biological siblings (n=26) during 500-ms binaural click trains (27- and 40-Hz) in an active oddball task.

**Results:** No group differences were observed in frequency-following responses (FFR) to 27- or 40-Hz stimuli, although higher 40-Hz power was associated with older age and better behavioral performance in ASD. The broad-band response from 180-250 ms was reduced in ASD for both stimulation frequencies—particularly in the low-frequency (<8 Hz) range—and significantly correlated with IQ and age. Siblings showed intermediate broad-band responses.

**Discussion:** While FFRs appeared intact in ASD, we observed reduced broad-band response in the transition period to the steady state FFR, which was specific to low (<8-Hz) frequencies—potentially reflecting reduced synchronization at timescales that correspond with slower, syllabic rhythms (∼4-8 Hz) occurring in natural speech. Intermediate responses in first-degree relatives suggest that this is related to genetic vulnerability for ASD and highlights its clinical relevance. These findings suggest intact sensory processing in ASD alongside possible top-down auditory feedback deficits, which may serve as heritable neurophysiological markers.

**Lay Abstract:** Children with autism may process sounds differently, which could contribute to challenges with attention and communication. Here, electroencephalography (EEG) measured how the brain responds to rapidly repeating sounds and found that, while basic sound processing was intact, children with autism showed significantly reduced low-frequency responses that may reflect difficulty tracking speech rhythm. Interestingly, unaffected siblings showed an intermediate response, suggesting this may be a heritable marker of neural differences in autism.

## Introduction

Autism spectrum disorder (ASD) is a neurodevelopmental condition characterized by social impairments, restricted, repetitive behaviors, and frequent sensory atypicalities [1]. Among these, altered auditory processing is well-documented in ASD [2-14], but its neurobiological basis and functional significance remain poorly understood. Electroencephalographic (EEG) studies—which offer millisecond-level temporal resolution [15, 16] of brain activity—consistently reveal smaller and/or slightly delayed auditory evoked potentials (AEPs) 50–200 ms post stimulus onset in comparison to age-matched neurotypical (NT) controls [3-14]. These auditory differences have been linked to heightened sensory sensitivity and reduced sound tolerance [17, 18], and may contribute to broader social and attentional difficulties observed in ASD.

Mounting evidence suggests that altered sensory processing in ASD may reflect a potential imbalance between excitatory and inhibitory (E/I) neural activity [19, 20]—an idea consistent with many animal models of autism [21-23] but for which there is only partial support from human studies [24, 25]. An especially useful tool to simultaneously probe both auditory processing and E/I circuitry is the auditory steady-state response (ASSR)— a sinusoidal frequency-following response (FFR) to rhythmic auditory stimulation, with major cortical generators in primary auditory cortex [26]. The 40-Hz ASSR, in particular, is thought to depend on the synchronized activity of GABAergic interneurons within the ventral division of the medial geniculate body (MGBv) [27], making it a sensitive marker of thalamocortical and cortical E/I dynamics. Beyond power of the FFR, the broad-band evoked response provides complementary insight into auditory processing [28-30]. Both the broad-band evoked response and ASSR power typically demonstrate right hemispheric dominance [31] and a positive association with age [32], making them informative benchmarks for studying auditory development in clinical populations [33].

Findings from ASSR studies in ASD have been mixed (**Supplementary Table 1**); while some show reduced ASSR power in ASD [34-37], others find no group differences [38-40]. To date, only two studies have examined the broad-band response to 40-Hz stimulation in ASD, with one finding a reduced broad-band response in the 150-250 ms period leading into the steady state ASSR [41], whereas the other—which focused on a wider window of 200-1000 ms—did not [37].

Our study was designed to clarify the conflicting evidence surrounding ASSR and the broad-band evoked response in ASD. We examined ASSR in a large cohort of children aged 8 to 12 years with ASD and in age- and IQ-matched controls—with and without a first-degree relative with autism—using highly accessible EEG techniques. Several features of our study enhance interpretability and novelty of ASSR findings in ASD. First, we restricted the age range to limit developmental variability. Second, whereas all prior ASSR studies in ASD used passive stimulation, we employed an active oddball detection task to monitor attentional engagement during EEG recording—a critical factor given the influence of attention on ASSR amplitude [42]. Third, we analyzed responses across multiple frequency bands; this decision was motivated by prior findings that not all children in this age range (8–12 years) exhibit a robust 40-Hz FFR [40, 41]. To address this, we included a lower-frequency (27-Hz) click-train stimulation condition, which may elicit more reliable responses in pre-adolescents [38]. Fourth, we evaluated group differences in the broad-band evoked response to help resolve mixed findings in the literature [37, 41]. Finally, we included unaffected first-degree relatives to assess whether ASSR alterations might reflect an endophenotypic trait [43, 44]. Although one prior study in ASD supports the potential of ASSR as a heritable endophenotype [34, 45], this possibility remains largely understudied.

Given substantial evidence of altered auditory processing and E/I imbalance in ASD—which may contribute to core symptoms—we hypothesized that individuals with ASD would show reduced ASSR power and altered broad-band responses. We further expected these neural measures to relate to clinical and cognitive features of autism, and that siblings would exhibit intermediate patterns consistent with a heritable neural signature.

## Methods

### Participants

Data from 55 children with ASD, 36 age-matched TD children, and 27 unaffected first-degree biological siblings of individuals with autism were originally included in the study. Among the initial participants, one TD participant, two participants with ASD, and one sibling were excluded from further analysis due to noisy EEG data (>15% bad channels or >50% bad epochs). Subsequent analyses included a final sample size of 114 (ASD: n=53, TD: n=35, sibling: n=26), as described in **Table 1**.

**Table 1.** Participant demographics. Continuous variables: mean ± standard deviation. Categorical variables: n (%). * Significant difference between TD and ASD (p-adj<0.05). ‡ Significant difference between ASD and siblings (p-adj<0.05). Missing clinical data - ADOS (ASD: 9); SRS-2 (ASD: 2, siblings: 3); FSIQ (ASD: 2, siblings: 1) *ASD – autism spectrum disorder; TD – typically-developing; ADOS-2 – Autism Diagnostic Observation Schedule, 2*^*nd*^ *edition; SRS-2 – Social Responsiveness Scale, 2*^*nd*^ *edition; IQ – intelligence quotient*

|  | ASD<br>(n=53) | TD<br>(n=35) | Siblings<br>(n=26) |
| --- | --- | --- | --- |
| **Sex, female \* †** | 10 (18.9) | 18 (51.4) | 15 (57.69) |
| **Age, years** | 10.84 ± 1.5 | 10.4 ± 1.8 | 10.56 ± 1.4 |
| **ADOS-2 Total** | 7.86 ± 2.1 | - | - |
| **SRS-2 Total T-score \* †** | 73 ± 12.2 | 46.89 ± 6.7 | 47.65 ± 7.7 |
| **Full Scale IQ (WASI-II or WISC-V) \*** | 95.04 ± 16.4 | 104.4 ± 16.1 | 103 ± 15.6 |
| **Handedness** |  |  |  |
| Left | 4 (7.5) | 2 (5.7) | 2 (7.7) |
| Ambidextrous | 1 (1.9) | 1 (2.86) | 0 (0) |

Participants ranged in age from 8 to 12 years old and were recruited without regard to sex, race, or ethnicity; TD and sibling group participants were recruited to match the ASD group by age. Intelligence quotients for verbal (V-IQ), and full-scale (FS-IQ) intelligence were assessed for all participants, using the Wechsler Abbreviated Scales of Intelligence (WASI-II; [46]) and Wechsler Intelligence Scale for Children (WISC-V; [47]). There were no significant group differences in full-scale IQ, but V-IQ was significantly higher for the TD group (vs. ASD). The Social Responsiveness Score (SRS-2; [48]) questionnaire was collected from all participants to obtain continuous measures of ASD characteristics related to social impairment.

To be included in the ASD group, participants had to meet diagnostic criteria for ASD on the basis of the following measures: *1*) autism diagnostic observation schedule 2 (ADOS-2) [49]; *2*) diagnostic criteria for autistic disorder from the *Diagnostic and Statistical Manual of Mental Disorders* (DSM-5); *3*) clinical impression of a licensed clinician with extensive experience in diagnosis and evaluation of children with ASD. Due to precautions during the COVID-19 pandemic, a subset of ASD participants (n=9) were not able to complete the ADOS-2 due to masking requirements. They instead underwent the Childhood Autism Rating Scale 2 (CARS-2) and Autism Diagnostic Interview-Revised (ADI-R) for diagnostic assessment. To be included in the typically-developing (TD) group, participants had to have no history of neurological, developmental, or psychiatric disorders, have no first-degree relatives with a diagnosis of ASD, and be in age-appropriate grade at school. To be in the sibling (SIB) group, the criteria were the same as for the TD group, plus having a sibling with a diagnosis of autism. Exclusionary criteria for all groups included: (1) known genetic syndrome (including syndromic cases of ASD), (2) history of or currently take medication for seizures in the prior 2-years, (3) exclusionary physical limitations (e.g. significant vision/hearing deficits), (4) were born prematurely (<35 weeks) and/or experienced prenatal/perinatal complications, or 5) FS-IQ <80.

All participants assented to the procedures and their parent or guardian signed an informed consent approved by the Institutional Review Board of the Albert Einstein College of Medicine. Participants received nominal recompense for their participation (at $15 per hour).

### Task

Auditory stimuli were 500-ms binaural click trains at either 27- or 40-Hz, presented through HD 650 Sennheiser headphones at 60 dB SPL. Inter-stimulus interval was randomly jittered between 488-788 ms. On 15% of trials, an oddball stimulus presented at a different frequency (27-Hz for 40-Hz trials, 40-Hz for 27-Hz trials) was randomly intermixed among the standards. Participants were instructed to respond via button-press when they identified an oddball stimulus, to promote attention to the auditory stimuli. Stimuli were presented in four randomly presented blocks of 100 trials—blocked by stimulus type (40-Hz standard, 27-Hz standard), consisting of 170 trials per standard frequency and 30 trials per oddball stimulus. Data collection for this paradigm took ∼10 minutes. Subsequent analyses include only the standard trials.

There were no significant differences in the final number of trials after epoch rejection across any groups for either condition (40-Hz – ASD: 155.25 ±19.54; TD: 153.51 ± 20.76; siblings: 155.12 ± 21.65; 27-Hz – ASD: 156.81 ± 22.49; TD: 159.49 ± 20.22; siblings: 157.23 ± 18.73).

### Data Collection

EEG recordings were collected from 70 active channels (10–20 system; 64 scalp channels and 6 external electrodes: 2 upper mastoids; 2 lower mastoids; 2 vertical Electrooculography (EOG) at a digitization rate of 512 Hz, using Active Two amplifiers (BioSemi, Amsterdam, The Netherlands) with an anti-aliasing filter (−3 dB at 3.6 kHz). Analog TTL triggers indicating stimulus onset and button presses were sent to the acquisition PC via Presentation (Neurobehavioralsystems) and stored digitally at a sampling rate of 512 Hz in a separate channel of the EEG data file. The system uses active electrodes and a direct current (DC) coupling with a hardware bandpass filter from DC to 150 Hz to reduce low-frequency drift and high-frequency noise. The recording reference was the Common Mode Sense (CMS) active electrode, which, together with the Driven Right Leg (DRL) passive electrode, forms a feedback loop to stabilize the reference signal and reduce common-mode noise. This referencing convention in BioSemi systems differs from traditional reference electrodes by dynamically compensating for potential differences between the scalp and the amplifier ground, effectively creating a “zero potential” point.

### Pre-Processing

EEG was analyzed using Python (MNE; [50]) and custom in-house scripts. Raw data were low-pass FIR-filtered at 80 Hz and high-pass FIR-filtered at 0.01 Hz, with a 0.01 Hz lower transition bandwidth and a 20 Hz upper transition bandwidth, corresponding to –6 dB cutoff frequencies at 0.01 Hz and 90 Hz, respectively.

Bad channels were detected automatically *(pyPrep*.*NoisyChannels; [51])* based on a combination of classical bad-channel detecting functions, which include: low signal-to-noise ratio, low RANSAC [52] prediction or correlation with other channels, abnormal amplitude or amounts of high-frequency noise, and near-flat channels. Subjects were excluded if >15% of channels were identified as bad; otherwise, bad channels were interpolated using the spherical-spline method [53].

Epochs were constructed from a -200 to +800 ms time window surrounding the onset of the stimulus. Independent Component Analysis (ICA) using the *fastica* algorithm (max 1,000 iterations) was applied to epoched data high-pass filtered at 1 Hz to identify and manually remove components corresponding to eye movements, including horizontal saccades and blinks.

Epochs were then baseline-corrected by subtracting the mean baseline per-individual from -200 ms pre-stimulus to stimulus onset. Epochs containing artifacts were excluded by automatic artifact rejection *(autoreject; [54])*, which utilizes a cross-validation metric to identify an optimal amplitude threshold [54]. Subjects were excluded if >50% of trials were rejected across all conditions.

Epochs were averaged at the individual subject level, for each of the stimulus conditions of interest, for subsequent analyses.

Finally, data was re-referenced to the average temporal electrodes (TP7, TP8) over the mastoid for subsequent statistical analyses. This approach is a commonly used reference technique for evaluating auditory event-related potentials, designed to optimize the measurement of the auditory cortical response [55, 56].

### Time-Frequency Analysis

Average power spectra were computed per-individual using complex Morlet wavelets, with a frequency resolution from 2 Hz to 45 Hz in 400 steps. The full-width at half-maximum (FWHM) of the wavelet ranged from 187ms to 162ms, corresponding to a spectral FWHM range of 1.5 Hz to 4 Hz. The temporal and spectral FWHM at 27 and 40-Hz were 167ms/3.5 Hz and 166ms/3.75 Hz, respectively. Time-frequency plots represent percent-change in power relative to a per-individual baseline of -150 to -50ms pre stimulus onset.

ASSR power was computed for the 200-500ms post-stimulus onset interval, following standard practice in previous studies [73, 74], and was extracted from fronto-central channels (FCz, FC3, FC4), which have been previously identified as representing the strongest responses to auditory click trains. Power was calculated for a 10-Hz range centered on the stimulation frequency (40-Hz condition: 35-45 Hz; 27-Hz condition: 22-32 Hz).

Using the same Morlet wavelet function in MNE, phase values were extracted as complex numbers per-individual at fronto-central channels (FCz, FC3, FC4). Complex power values were cropped to a 1-Hz range around the target frequency (39.5-40.5 Hz and 26.5-27.5 Hz) and subsequently converted to phase angles in radians. These phase angles were re-expressed at each time point as the difference from the expected angle [TD mean]. The mean difference per-individual within a given time window was defined as the phase-locking angle (PLA). PLA, initially expressed as difference in radians, was converted to Z-scores by subtracting the TD mean and dividing by standard deviation of the TD group, allowing us to perform linear—rather than circular (i.e. Watson-Williams test)—statistics.

### Statistical Analysis

Group differences (ASD, TD, SIB) in neural (Z-scored PLA), clinical (age, IQ, SRS-2, ADOS-2), and behavioral (reaction time, percent correct, percent false alarms, detection sensitivity) measures were assessed by ANOVA testing using R (v.4.4.0) with α=0.05. Data were assessed for normality and homogeneity by Shapiro-Wilk test [57] and Levene’s test [58], respectively. Parametric data was assessed primarily by ANOVA, with follow-up post-hoc testing (equal variance: Tukey’s [59]; unequal variance: Games-Howell [60]). The Kruskal-Wallis test [61] with post-hoc Wilcoxon was used to evaluate non-parametric data, with Benjamini-Hochberg [62] correction for multiple tests. Analysis of categorical variables (biological sex) were assessed by chi-squared or fisher’s (n<10) test [63] with Bonferroni-correction [64]for multiple tests.

Statistical analysis of neural measures (ASSR power and amplitude of the broad-band ERP) that have previously demonstrated hemispheric and age-related differences [32, 38, 39, 41], was conducted using a linear mixed-effects model (LMM; [65]). This approach allowed for the incorporation of channel as a variable (to consider lateralization of the response) while assessing group differences in non-parametrically distributed data. Follow-up pairwise comparisons of the LMM were obtained by estimated marginal means (emmeans; [66]) to compute contrasts between Group x Hemisphere (or channel for ASSR), with Age as a covariate, applying a Tukey correction. Correlations between clinical (FS-IQ, age, ADOS-2, SRS-2), behavioral (detection sensitivity, percent correct, percent false alarms, mean reaction time) and neural (broad-band, ASSR power, and phase delay) measures were evaluated by either Pearson [67] or Spearman [68] tests, as determined by normality (assessed by Shapiro-Wilk test).

Statistical analysis of the broad-band signal was conducted across three time windows (0–180 ms, 180–250 ms, 250–500 ms), defined based on the shape of the broad-band response (see **Fig. 2**). PLA analysis was performed on the second two of these windows (the broadband slope leading into the steady-state response (180–250 ms) and (ii) the steady-state response itself (250–500 ms) to assess potential phase delay in ASD or SIBS compared to controls. ASSR power was evaluated within the peak steady-state window (200–500 ms), following standard practice in previous studies [69, 70].

To ensure comparability with existing literature while also examining the impact of attention, we analyze the data in two ways: (1) including all participants, to align with prior studies (presented first), and (2) excluding behavioral outliers, allowing for more control of attentional effects on ASSR.

## Results

### ASSR Power

Time-frequency plots show marked increase in power compared to baseline at the stimulated frequency with striking similarity across all three groups, for both 40- (**Fig. 1a**) and 27-Hz (**Fig. 1c**) conditions. Examination of the scalp topography of ASSR power in the 200-500 ms window of interest suggested similar scalp distributions over fronto-central scalp across the groups for both the 40-Hz and 27-Hz conditions, with smaller response to 27-Hz compared to 40-Hz auditory stimulation (see **Fig. 1a-b** and **Fig. 1c-d**). ASSR power did not show a significant effect of Group for either 40-Hz (**Fig. 1a-b**) or 27-Hz (**Fig. 1c-d**) conditions. We identified a significant main effect of Channel in both 40-Hz (F=5.426, p=0.005) and 27-Hz (F=5.264, p=0.006) conditions suggestive of a right hemisphere bias in the response. Follow-up post-hoc analysis in the 40-Hz condition suggests that left fronto-central ASSR power is significantly lower relative to midline (FC3-FCz: β=-0.05, p=0.004). In the 27-Hz condition, we identified significantly greater ASSR power over right relative to left fronto-central scalp (FC3-FC4: β=-0.035, p=0.006). Moreover, we identified a significant main effect of Age in the 40-Hz condition (F= 13.4, p <0.001).

**Figure 1.**
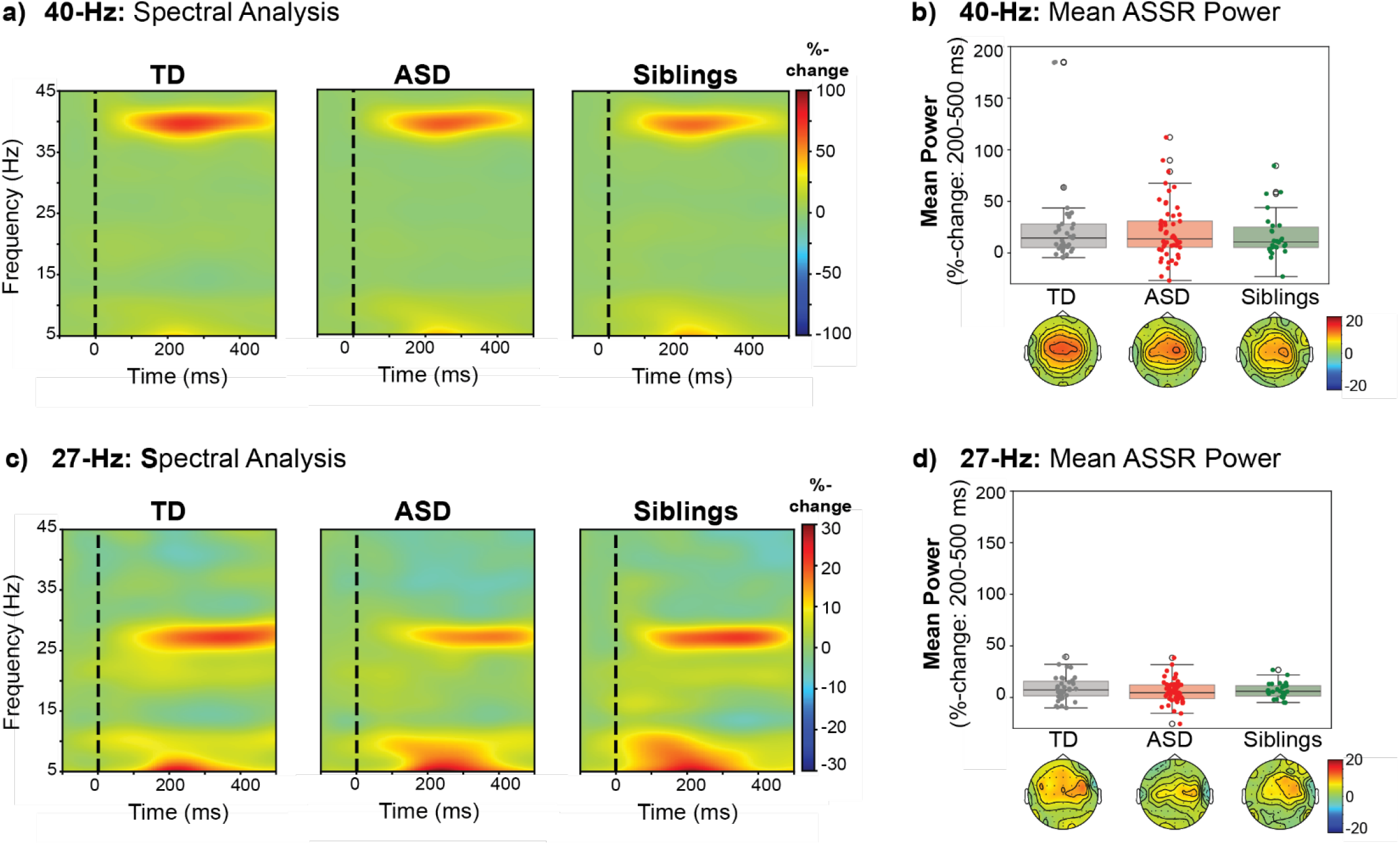
Similarity in 40 and 27-Hz ASSR Power across groups. **a, c)**Time-frequency plots display Morlet-transformed power, expressed as %-change relative to baseline, over fronto-central scalp (averaged over FCz, FC3, FC4), for **(a)** 40-Hz and **(c)** 27-Hz conditions, separated by group. **b, d)** Mean ASSR power (averaged power within 5-Hz of the frequency following response from 200 to 500ms) is shown for each group (TD: grey; ASD: red; siblings: green) for **(b)** 40-Hz and **(d)** 27-Hz condition. Topographical maps illustrate the scalp distribution of ASSR power from 200 to 500 ms for each group, plotted as %-baseline.

### Phase Locking Angle (PLA)

To further compare the timing of the FFR across groups, we evaluated the phase-locking angle (PLA) of siblings and ASD, referenced to the TD mean. The PLA difference per individual was calculated at each time point by subtracting the respective TD phase angle (at 40Hz and 27Hz for the respective stimulation conditions), which was then averaged across the time windows of interest (180-250 ms; 250-500 ms).

Comparison of z-scored PLA (**Supplementary Figure 1**) did not reveal significant differences from TD in the ASD or sibling group, indicating intact frequency following neuronal responses for both conditions.

### Broad-Band Evoked Response

The ERP scalp topographic maps and waveforms for 40-Hz (**Fig. 2a)** and 27-Hz (**Fig. 2c)** conditions reveal similar responses across the groups, which are comprised of two positive-going deflections over fronto-central scalp in the first 180 ms of the response, and a downward sloping response from ∼180 to 250ms with bilateral fronto-central foci that transitions into the steady-state frequency-following response (250-500 ms) over fronto-central scalp.

Findings from the LMM failed to show significant differences between Group or Channel in the 0-180 ms time window, although there was a significant main effect of Age in both 27-Hz (F=5.462, p=0.012) and 40-Hz (F=12.784, p=0.001) conditions.

**Fig 2.**
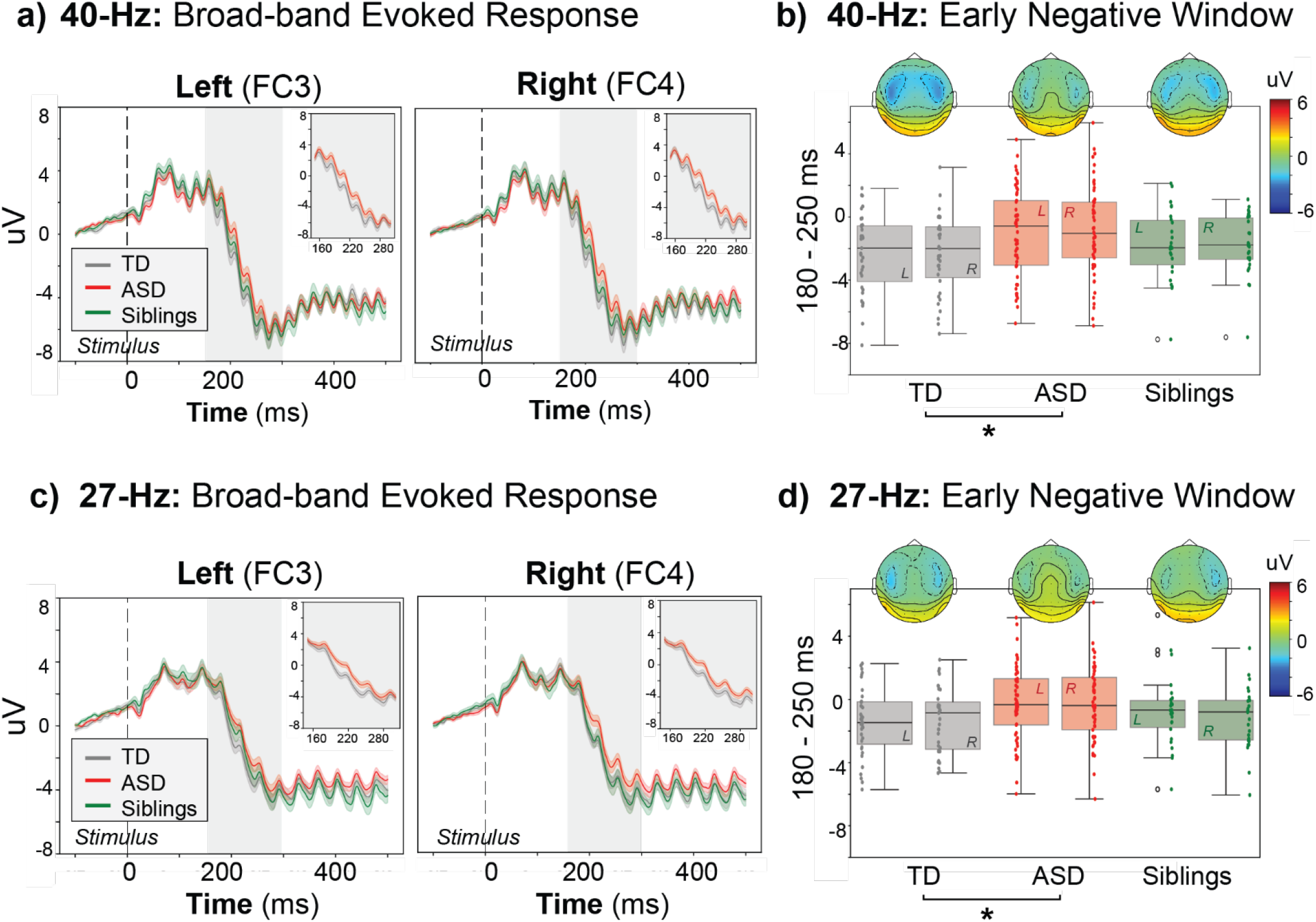
Significant Group Differences in 180-250 ms Window of the 40 and 27-Hz Broad-Band Evoked Response. **a, c)** broad-band evoked response, separated by group (TD: grey, ASD: red, siblings: green). To better visualize group differences in the 180-250 ms window, grey boxes zoom in to the TD (grey) and ASD (red) evoked response during this time. **b, d)** boxplots of mean amplitude in early-negative-going time window (180-250 ms) per-individual, separated by group (TD: grey, ASD: red, siblings: green) and hemisphere (L: left, R; right) and hemisphere (L: left, R; right). Topographies show the scalp distribution of broad-band evoked response in the 180-250 ms window, separated by group. Asterisks demonstrate significant main effects of Group (p-val <0.05).

For the 180-250 ms window (**Fig. 2b, 2d)**, there was a significant main effect of Group in both 27-Hz (F=3.42, p=0.036) and 40-Hz (F=3.926, p=0.023) conditions. Follow-up post-hoc testing demonstrated a significant difference between ASD and TD groups in the 40-Hz (β=-1.491, p=0.019) and 27-Hz (β=-1.208, p=0.032) conditions, without differences between TD vs. siblings or ASD vs. siblings.

In the 40-Hz condition only, analysis of the 250-500 ms window revealed a significant interaction between Group and Hemisphere (F=3.258, p=0.042). Follow-up post hoc analysis revealed a marginal, but statistically non-significant, difference between the left (FC3) and right (FC4) hemispheres in both the TD (FC3-FC4: β=0.306, p=0.055) and ASD (FC3-FC4: β=-0.202, p=0.118) groups.

### Exploratory follow-up of the group difference in the 180-250ms window

The modulation in the 180-250 ms window appeared to represent a latency shift in the onset of the negative going response in lower frequency EEG activity. To further visualize potential drivers of the group differences identified in the ERP from 180-250 ms, band-filtering of the response at commonly studied frequency bands was performed (delta: 0.5-4Hz, theta: 4-8 Hz, alpha: 8-12 Hz, low beta: 12-20 Hz). Visually, delta and theta bands (shown in **Supplementary Figure 2**) appeared to drive the underlying shape of the broad-band evoked response shown in **Fig. 1**. Subsequently, we filtered the broad-band to just these low-frequencies (delta and theta bands; 0.5-8 Hz). As demonstrated in **Fig. 3**, this low-pass filtered-ERP closely mirrors the shape of the original broad-band response (**Fig. 1**). Furthermore, the low-frequency evoked response topography (**Supplementary Figure 3**) showed foci over fronto-central scalp during the negative-going response (180-250 ms) of both conditions, suggesting the group differences identified during this period are driven by lower frequency neural activity within 0.5-8 Hz.

**Fig. 3.**
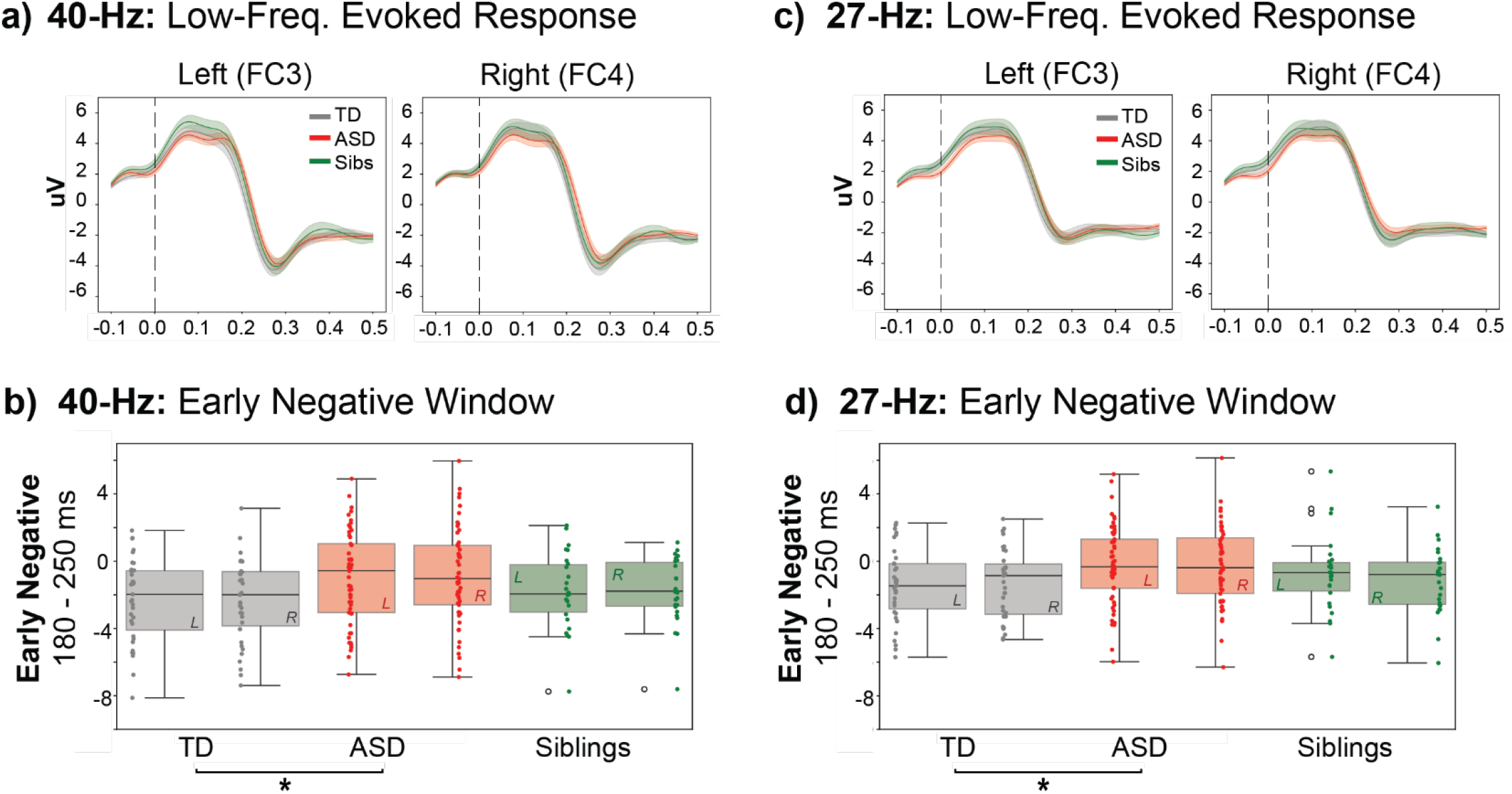
Significant Group Differences in Amplitude of the 40 and 27-Hz Low-Frequency (0.5-8 Hz) Evoked Response. **a, c)** low-frequency filtered (0.5-8 Hz) evoked response, separated by group (TD: grey, ASD: red, siblings: green). **b, d)** boxplots of mean amplitude in early-negative-going time window (180-250 ms) per-individual, separated by group (TD: grey, ASD: red, siblings: green) and hemisphere (L: left, R; right). Asterisks demonstrate significant main effects of Group (p-val <0.05).

A LMM on the 0.5-8 Hz data within this time window (180-250 ms) revealed a main effect of Group in both the 40-Hz (F=4.012, p=0.021) and 27-Hz (F=3.481, p=0.034) conditions. Post-hoc comparison demonstrated a significant difference in amplitude between TD and ASD in both the 40-Hz (β=-1.058, p=0.03) and 27-Hz (β=-1.312, p=0.018) conditions. Siblings did not significantly differ from either the TD or ASD groups in either condition or hemisphere. This suggests that the previously identified broad-band group differences are largely driven by frequencies below 8 Hz, as demonstrated in prior work [41].

### Behavioral Results, and Task Performance and the ASSR

Attention has been shown to influence the amplitude of the ASSR. Differences in how attention is allocated may account in part for inconsistent findings regarding group differences in the literature. In the present study, participants performed an oddball detection task on the eliciting stimuli in which they were to respond to the suprathreshold deviant tones as quickly as possible. Performance metrics allowed us to assess how well they were attending the click trains.

Statistical analysis revealed a main effect of Group in both mean reaction time (**Fig. 4A**; F =3.559; p-val=0.032) and percentage of false alarms (**Fig. 4D**; χ^2^=11.239; p-val=0.004). Follow-up post-hoc analysis (Tukey) of mean reaction time demonstrated significantly faster response times in the ASD vs. TD group (TD-ASD mean difference=56.601 ms; p-val=0.024). Furthermore, pairwise comparison (Wilcoxon-ranked sum) showed the ASD group had significantly higher percentage of false alarms than the TD group (W=1,287; p-val=0.003). Post-hoc analysis for differences in both reaction time and false alarms was insignificant between siblings and either TD or ASD groups.

**Fig 4.**
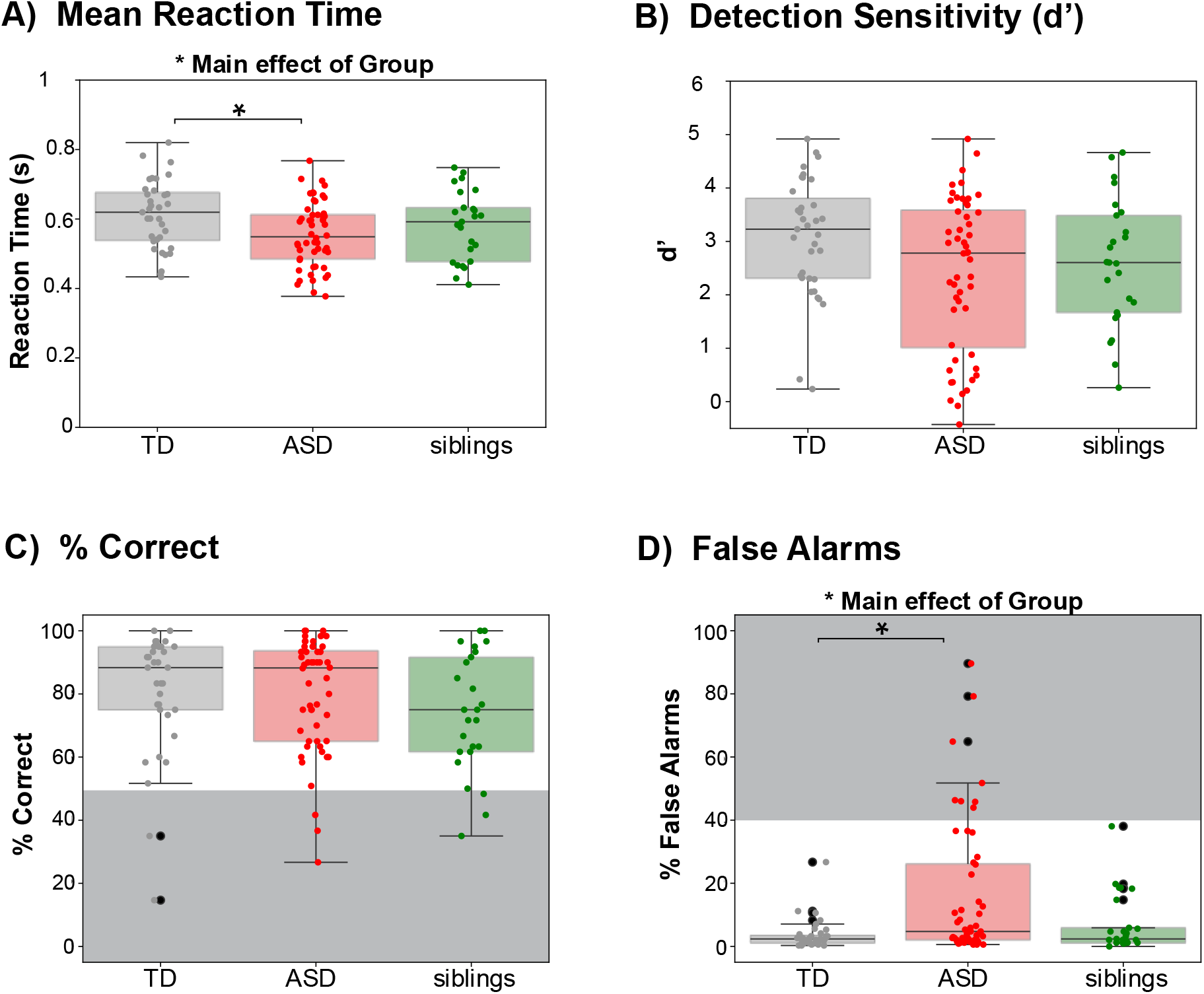
Group Differences in Oddball Discrimination. Boxplots of **A)** detection sensitivity (d’), which refers to the difference between Z(hit rate) and Z(false alarm rate), per-participant, **B)** mean reaction time of correct responses, **C)** percent correct, and **D)** percent false alarms (responses to a non-deviant tone), separated by group (TD: grey, ASD: red, siblings: green). Grey boxes demark thresholds for behavioral outlier exclusions (<50% correct and/or >40% false alarms). Participants with 0% correct responses were excluded from plots and statistical analysis.

To understand if attentional variability influenced the ASSR data, poor behavioral performers—defined as having <50% correct responses and/or >50% false alarms—were excluded from the analysis. Figure 4C-D illustrates how poor performers were distributed across the groups. Notably, they were far more prevalent in the ASD group (23%; N=12), than either the TD (5.7%; N=2) or the sibling (15.4%; N=4) groups. Since all participants could easily detect the oddball stimulus, these individuals were either a) not participating in the task (evidenced by few correct responses) or b) responding to the stimuli indiscriminately (evidenced by many false alarms).

Re-analysis of behavioral data after removal of these participants demonstrated a significant effect of Group for mean reaction time (F=3.95; p-val=0.023). Pairwise comparison revealed that the ASD group was still significantly faster than the TD group in response time (TD-ASD mean difference=61.365ms; p-val=0.016), although error rates no longer significantly differed.

### Analysis of Electrophysiological Data After Removal of Poor Performers ASSR Power

As with the full dataset, analyses did not identify group differences in ASSR power with either stimulation frequency. There was a significant main effect of Age in the 40-Hz condition (F=7.196, p=0.009) and a significant main effect of Channel in both 40-Hz (F=4.837, p=0.009) and 27-Hz (F= 6.197, p=0.002) conditions, suggestive of a right hemisphere bias in the response. Follow-up post-hoc analysis in the 40-Hz condition suggested that left fronto-central ASSR power was significantly lower relative to midline (FC3-FCz: β=-0.053, p=0.006). For the 27-Hz condition, ASSR power was significantly greater over right relative to midline (FC4-FCz: β=0.03, p=0.034) and left fronto-central scalp (FC3-FC4: β=-0.041, p=0.002).

### Broad Band Response

Analysis of the 180-250 ms window (**Fig. 5b, 5d)** revealed a significant main effect of Group in the 27-Hz condition (F=3.202, p=0.045) due to a more negative-amplitude response in TD relative to ASD (β=-1.206, p=0.045). The main effect of Group was marginal in the 40-Hz condition (F=2.797, p=.066). There were no significant differences between TD vs. siblings or ASD vs. siblings.

**Fig 5.**
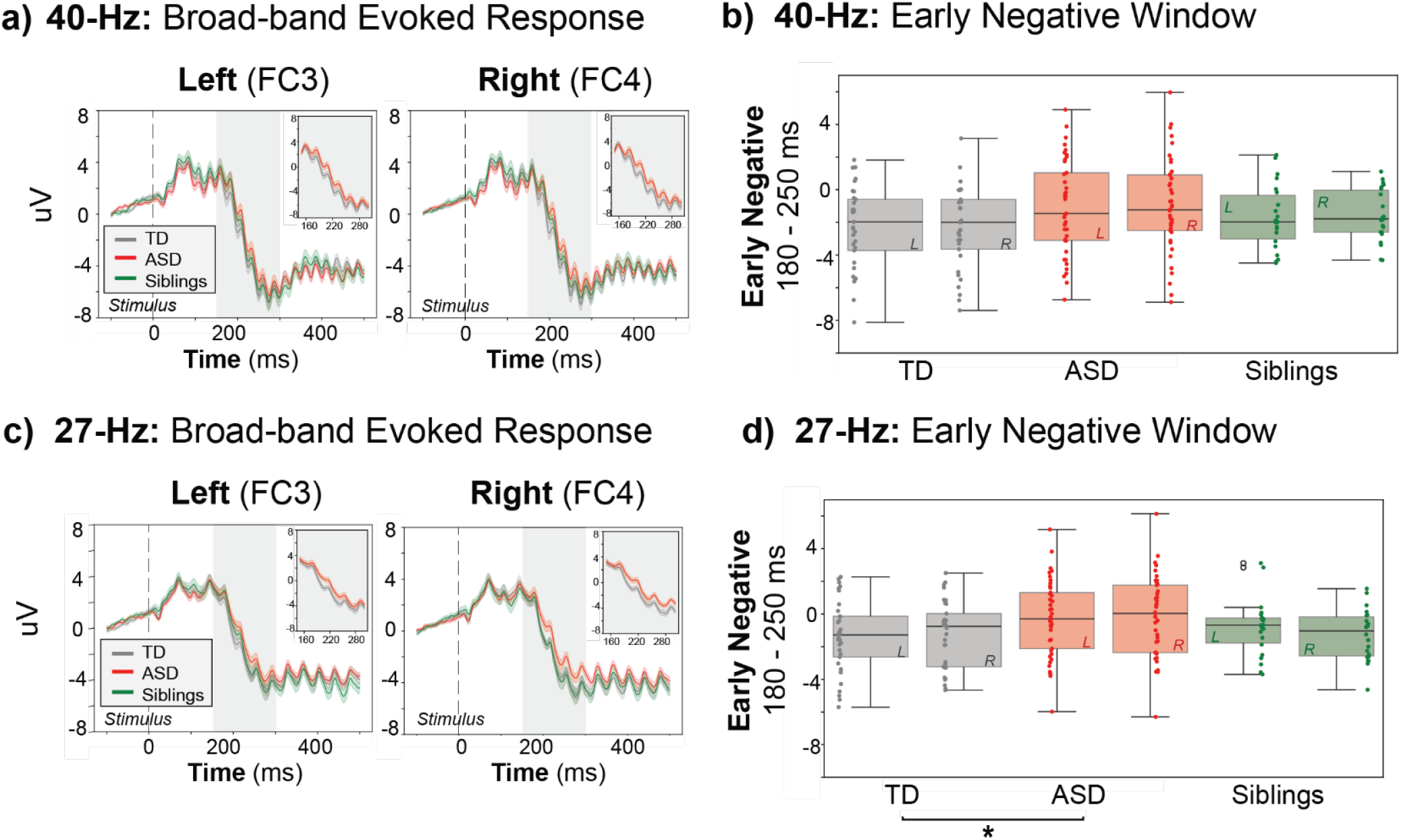
40 and 27-Hz Broad-Band Evoked Response After Removal of Poor Performers. **a, c)** broad-band evoked response, separated by group (TD: grey, ASD: red, siblings:). **b, d)** boxplots of mean amplitude in early-negative-going time window (180-250 ms) per-individual, separated by group (TD: grey, ASD: red, siblings: green) and hemisphere (L: left, R; right). Asterisks demonstrate significant main effects of Group (p-val <0.05).

Testing of the steady-state (250-500 ms) portion of the broad-band waveform revealed a significant Group x Hemisphere interaction in the 40-Hz condition (F=4.077, p=0.02). Follow-up post hoc analysis revealed a significant hemisphere effect (between FC3 and FC4) in the ASD group (β=-0.325, p=0.028), and a marginal, non-significant difference in the TD group (β=0.297, p=0.071).

### Clinical Correlates

Using the full dataset, possible correlations between neural measures (broad-band evoked response, ASSR power) and clinical (IQ, ADOS-2, SRS-2, age) or behavioral (detection sensitivity, mean reaction time, % correct, % false alarms) variables of interest (**Fig. 6**) were probed.

**Fig 6.**
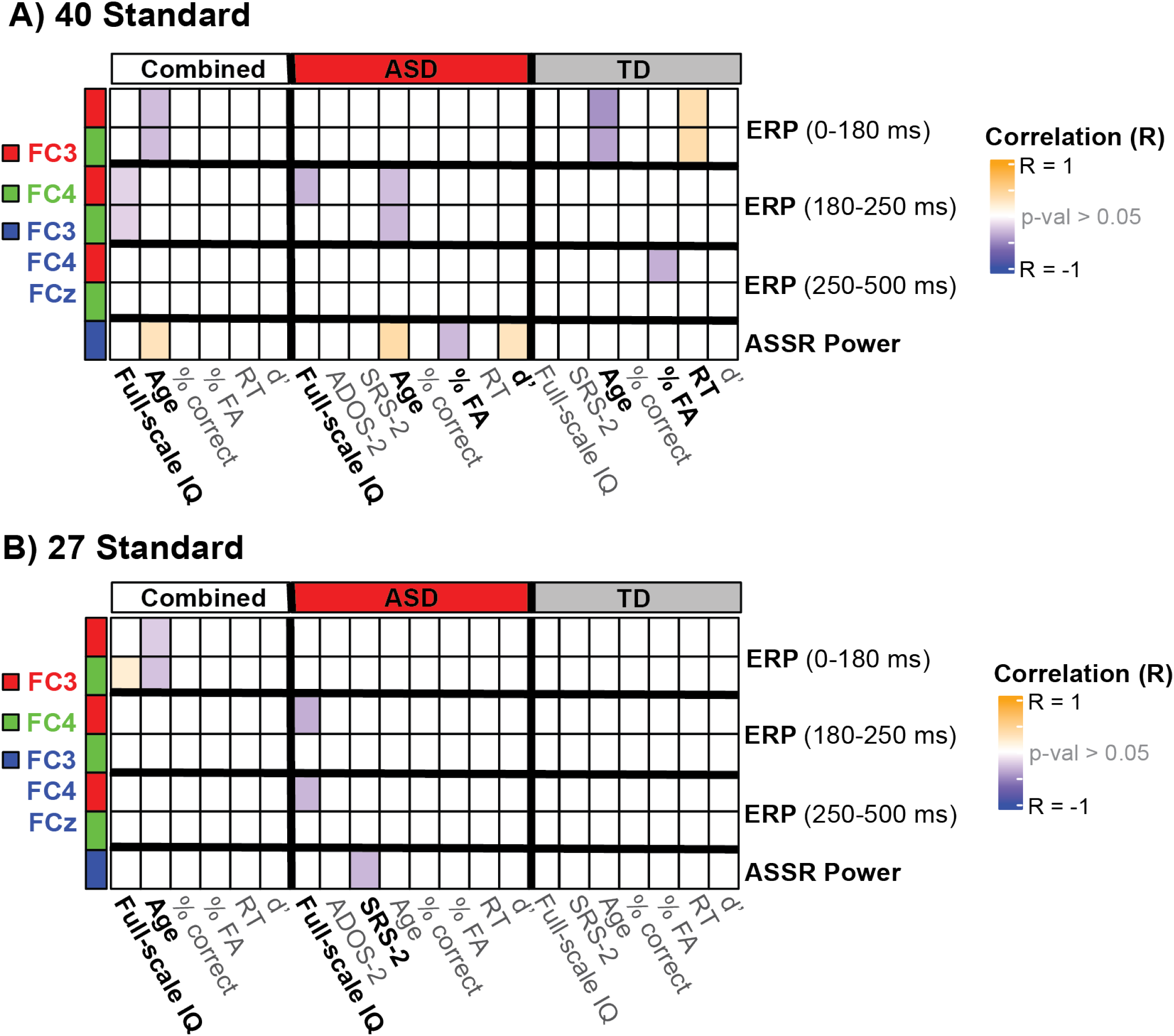
40 and 27-Hz Clinical, Behavioral and Neural Correlates. Heatmaps for 40-Hz **(A)** and 27-Hz **(B)** clinical correlations of neural measures (early-positive [0-180 ms], early-negative [180-250 ms] broad-band ERP, steady-state [250-500 ms] broad-band ERP, ASSR power). Non-significant correlations (p-val <0.05) are shown in white. Significant correlations (p-val <0.05) are bolded and represented by color-scale denoting the R-value of the correlation (orange: positive; blue: negative). Combined group correlations include all participants (TD, ASD, siblings), including behavioral outliers. ASD and TD groups are also evaluated separately.

#### ASSR power

Greater 40-Hz ASSR power significantly correlated with older age (Combined: R=0.3, p=0.001; ASD: R=0.382, p=0.005), fewer false alarms (ASD: R=-0.298, p=0.03) and greater detection sensitivity (ASD: R=0.291, p=0.038). 27-Hz ASSR power negatively correlated with SRS-2 score (ASD: R=-0.309; p=0.027), indicating lower amplitude frequency-following response was associated with greater social impairment.

#### Broad-band (180-250 ms)

In the 40-Hz condition the broad-band response within the 180-250 ms time window negatively correlated with age (ASD-FC3: R=-0.273, p=0.048; ASD-FC4: R=-0.295, p=0.032) and IQ (Combined -FC3: R=-0.187, p=0.05; Combined-FC4: R=-0.191, p=0.045; ASD-FC3: R=-0.316, p=0.024), suggesting a more negative amplitude during this window is associated with older age and higher IQ. A similar clinical correlate was identified in the 27-Hz condition; a more negative amplitude in the early negative broad-band evoked response significantly correlated with higher IQ in the ASD group, but only on the left side (ASD-FC3: R=-0.348; p=0.012).

## Discussion

In this study, we investigated auditory frequency-following and broad-band evoked responses to two click-train stimulation frequencies (27 Hz and 40 Hz) in children with ASD, their unaffected siblings, and typically developing (TD) controls. Using a well-powered sample, we found no evidence for group differences in the magnitude (power) or timing (PLA) of the FFR at either stimulation frequency. Intact FFRs were evident whether-or-not attention was well-controlled between the groups (as assessed by meeting performance thresholds on the oddball task), suggesting that this is a robust response that was not greatly influenced by differences in attention. In contrast, the ASD group showed a reduction in the broad-band evoked response from 180–250 ms post-stimulus—particularly in the low-frequency range (<8 Hz). Unaffected siblings exhibited intermediate response amplitudes, suggesting a potential endophenotypic pattern.

### Intact ASSR & PLA in TD vs. ASD children

Despite significant efforts to optimize detection—such as controlling for attention and using a restricted age cohort—we did not find evidence of deficits in ASSR power or PLA for either stimulation frequency. Although findings in prior ASSR studies are mixed, our results align with several well-powered MEG studies in similar age ranges which report preserved ASSR power and synchronization in ASD [38-41]. In contrast, other ASD studies demonstrate hemispheric differences in ASSR power [34-37]—including Arutiunian et. al [37], which examined a similarly-aged pre-adolescent cohort. Such discrepancies may reflect differences in cohort characteristics (e.g., symptom profile or cognitive ability), as well as variations in stimulus properties or analytic methods. Importantly, no previous studies have monitored attentional engagement during the task. In our study—which incorporated an active oddball detection task to index attention—ASSR responses remained stable regardless of task performance, suggesting that the frequency-following response is robust to moderate attentional variability.

Given the well-established link between magnitude (power) and timing (PLA) of the gamma-band FFR and the function of fast-spiking parvalbumin-positive interneurons [71], these findings argue against widespread E/I imbalance in primary auditory cortex during this developmental window. While one could argue that 40-Hz ASSR may not be reliably evoked in this age group—given its positive correlation with age and GABAergic system maturation (both here and in prior works [32])—group-level responses were present, and differences would be expected if ASD were associated with a reduction or developmental shift in these processes. However, this does not preclude the possibility that E/I dysregulation contributes to ASD pathophysiology in other cortical regions, such as higher-order association areas, or under different cognitive demands. Previous studies employing magnetic resonance spectroscopy (MRS) do offer evidence supporting regionally specific alterations in E/I balance in ASD, showing reduced GABA (inhibitory) concentrations in the left auditory cortex [72]—observed in both individuals with ASD and their unaffected siblings—and elevated GABA alongside reduced glutamate (excitatory) in the right temporal cortex, with the latter correlating with greater autism symptom severity [73]. It is also possible that individuals with ASD who fall outside the developmental or cognitive profile represented in this cohort—particularly those with more severe symptoms or co-occurring conditions—may exhibit different patterns of ASSR and E/I imbalance.

The clinical significance of ASSR in ASD remains uncertain. While some prior studies suggest positive associations with language [37] and cognitive ability [39, 74], the broader functional implications of these differences are not well understood. Here, we identified a significant association between greater 40-Hz ASSR power and improved behavioral performance (higher detection sensitivity and lower false alarm rate); however, it is possible that this relationship was partially driven by age-related effects, as ASSR power increases with development and older participants generally demonstrate better behavioral performance. Future studies with larger samples are needed to disentangle age-related maturation from task-specific auditory processing efficiency.

### Broad-band evoked response amplitude differences in TD vs. ASD children

In contrast to the ASSR, the broad-band evoked response revealed clear group differences. Children with ASD exhibited a significant reduction in amplitude in the downward slope leading into the steady-state response (180-250 ms) in both 27-Hz and 40-Hz conditions. Time-frequency analyses confirmed that this reduction was specific to low-frequency activity (<8 Hz). To date, only two prior studies evaluate the broad-band response in ASD, yielding conflicting findings. Our results closely align with the MEG findings of Stroganova et. al [41], who reported a significantly delayed and attenuated response in the left hemisphere during the 150-250 ms time window—specific to low-frequencies (<9 Hz)—within the ASD group. Methodological differences—such as the use of monaural stimulation, broader age ranges, or variation in IQ—may account for the lateralization differences across studies. Importantly, the replication of this early evoked response alteration across independent, well-powered cohorts and recording modalities (MEG and EEG) suggests it is a robust neural signature of auditory processing differences in ASD. In contrast, Arutiunian et. al [37] failed to identify significant group differences in the broad-band evoked response, although they examined a much larger time window (200-1,000 ms). In the present study, we also failed to identify significant group differences in the extended steady-state broadband response (250–500 ms), suggesting that the observed broad-band delay is highly temporally constrained.

The functional role of the broad-band delay—particularly in low-frequencies—is not well described in previous literature. Theoretically, even small delays in early auditory cortical processing, especially within the 150–250 ms window preceding the steady-state response, could disrupt the timing of auditory information integration. Such temporal imprecision may impair the brain’s ability to process rapid streams of sensory input, like speech or socially relevant sounds, which depend on precise neural synchronization. Moreover, the broadband response is largely composed of low-frequency activity—including delta and theta bands—which have been implicated in word comprehension [75, 76] and entrainment to syllabic rate [75, 77, 78], respectively. Recent evidence also suggests that auditory selective attention enhances theta-range activity in the auditory nerve during sound anticipation, supporting the idea that the low-frequency broadband response may index top-down modulation of auditory processing [79]. As such, the reduced activity in the theta band that we observe may indicate impaired auditory feedback processing, though this remains speculative and requires further investigation. Future studies—which should likely include animal models of ASSR—should specifically evaluate the mechanisms underlying this stage of auditory processing. Additional work in humans may also incorporate studies of natural speech, in addition to simple auditory stimuli, to evaluate the functional link between the low-frequency delay in evoked responses and speech comprehension, given mounting evidence for atypical speech processing in ASD [80-82].

### Intermediate Broad-Band Response in Siblings

Here, we directly tested the hypothesis that ASSR responses may serve as an endophenotypic marker for ASD by measuring ASSR in unaffected siblings of individuals with a diagnosis of ASD. The present data did not reveal group differences in the FFR, indicating that this is not a robust marker of ASD. However, we did identify significant group differences in the 180-250 ms time window of the broad-band response. While the ASD group showed a significant reduction in amplitude relative to the TD group, the sibling group did not differ significantly from either but appeared intermediate—falling between ASD and TD participants. This pattern is consistent with an endophenotypic response reflecting genetic vulnerability to ASD. However, given the modest effect size and substantial overlap between groups, this relationship should be interpreted cautiously and warrants further investigation to establish sensitivity and specificity across broader samples.

### Effect of Behavior

To our knowledge, this is the first ASSR study in ASD to incorporate an active auditory oddball task, enabling us to monitor and evaluate how variability in attention might influence auditory evoked responses. Despite suprathreshold target presentation, the ASD group demonstrated markedly poorer behavioral performance relative to the TD group, consistent with generalized attentional differences reported in ASD [83, 84]. Removal of 18 participants due to poor behavioral performance, which makes up 15.8% of the original clinical cohort, influenced identification of group differences in the broad-band delay. Whereas we initially identified a main effect of Group in both 40-Hz and 27-Hz conditions, significant group differences were present only in the 27-Hz condition after behavioral outlier removal. This likely reflects loss of statistical power, in which small effects were no longer detectable at the group level with a markedly reduced sample size. Alternatively, it may reflect attentional modulation of the auditory evoked response. As shown in previous literature [85], enhanced evoked responses to attended (vs. unattended) auditory inputs could minimize group differences, although further work is needed to more directly test attentional modulations of ASSR in ASD, and if this is influenced by stimulation frequency.

## Conclusion

In sum, our findings suggest that auditory steady-state responses—both in terms of power and phase—are largely preserved in children with ASD, indicating intact local cortical entrainment and no strong evidence of widespread E/I imbalance in primary auditory regions during this developmental window. In contrast, the reduced broad-band evoked response observed in the ASD group—particularly in the low-frequency (<8 Hz) range— points to subtle alterations in early auditory processing that may reflect impairments in top-down modulation or temporal alignment of auditory processing. The intermediate responses seen in unaffected siblings further suggest that this neural signature may reflect a heritable marker of vulnerability. Together, these results highlight the value of combining ASSR and broad-band measures to better characterize auditory system function in ASD and underscore the importance of incorporating developmental context, task engagement, and familial risk when interpreting group-level neural differences.

## Author Contributions

J.J.F, S.M conceived the study. D.C, T.L implemented the experiment and recorded the data. M.D pre-processed and analyzed the data under the supervision of T.V and S.M. M.D wrote the first draft of the manuscript. J.J.F, S.M, and T.V edited the manuscript.

## List of Abbreviations

ADOS-2: Autism Diagnostic Observation Schedule, 2nd edition
ADI-R: Autism Diagnostic Interview–Revised
AEP: Auditory Evoked Potential
ANOVA: Analysis of Variance
ASD: Autism Spectrum Disorder
ASSR: Auditory Steady-State Response
CARS-2: Childhood Autism Rating Scale, 2nd edition
DSM-5: Diagnostic and Statistical Manual of Mental Disorders, 5th edition
d′: Detection sensitivity
EEG: Electroencephalography / Electroencephalogram
E/I: Excitatory/Inhibitory
EOG: Electrooculography
ERP: Event-Related Potential
FFR: Frequency-Following Response
FS-IQ: Full-Scale Intelligence Quotient
FWHM: Full Width at Half Maximum
GABA: Gamma-Aminobutyric Acid
Hz: Hertz
ICA: Independent Component Analysis
IQ: Intelligence Quotient
ITC: Inter-Trial Coherence
L: Left (hemisphere or electrode)
LMM: Linear Mixed-Effects Model
MGBv: Medial Geniculate Body, ventral division
MEG: Magnetoencephalography
MRS: Magnetic Resonance Spectroscopy
ms: Millisecond(s)
NT: Neurotypical
PLA: Phase-Locking Angle
R: Right (hemisphere or electrode)
SIB: Sibling group (unaffected biological siblings of individuals with ASD)
SNR: Signal-to-Noise Ratio
SRS-2: Social Responsiveness Scale, 2nd edition
TD: Typically Developing
V-IQ: Verbal Intelligence Quotient
WASI-II: Wechsler Abbreviated Scale of Intelligence, 2nd edition
WISC-V: Wechsler Intelligence Scale for Children, 5th edition

## Acknowledgements & Funding

This work was supported by a grant from the Simons Foundation Autism Research Initiative (SFARI Award # 874845, SM). Support for recruitment and phenotyping of participants was provided by the Human Clinical Phenotyping Core of the NICHD funded Rose F. Kennedy Intellectual and Developmental Disabilities Research Center (P50 HD105352). Work at the collaborating site in Rochester is supported through the B. Thomas Golisano Intellectual and Developmental Disabilities Research Institute (UR-IDDRC), which is supported by a center grant from the Eunice Kennedy Shriver National Institute of Child Health and Human Development (P50 HD103536 – to JJF). The content is solely the responsibility of the authors and does not necessarily represent the official views of the National Institutes of Health.

