## Supplementary Table 1 for "Testing the auditory steady-state response (ASSR) to 40-Hz and 27-Hz click trains in children with autism spectrum disorder and their first-degree biological relatives: A high-density electroencephalographic (EEG) study"

| *Paper* | *Design* | *Participants* | *Beta* | *Gamma* | *Other Findings* |
|---|---|---|---|---|---|
| Ahlfors, 2024 [1] | MEG: ASD vs. TD (25 & 43 Hz) | CHILDREN *ASD: n=22* *TD: n=31* **(6-17 YO)** | No difference (ASD vs. TD); dominance in R hemisphere (ASD, TD) | No difference (ASD vs. TD); dominance in R hemisphere (ASD, TD) | No difference (ASD vs. TD) in double-frequency auditory response |
| Arutiunian, 2023 [2] | MEG: ASD vs. TD (40-Hz) | CHILDREN *ASD: n=20* *TD: n=20* **(7-14 YO)** | - | **Reduced power & ITC in R hemisphere (ASD vs. TD);** no evoked response difference (ASD vs TD) | R-side ITC positively correlated with age (TD only); L-side ITC positively correlated with mean language score (ASD) |
| Stroganova, 2020 [3] | MEG: ASD vs. TD (40-Hz) | CHILDREN *TD: n=35* *ASD: n=35* **(7-12 YO)** | - | Dominance in R hemisphere (ASD, TD); **evoked response delayed and displaced in L hemisphere (ASD vs. TD)** | ASSR positively correlated with age (ASD, TD); no correlation between evoked response delay & intelligence or ASD severity |
| Seymour, 2020 [4] | MEG: ASD vs. TD (40-Hz) | ADOLESCENTS *ASD: n=18* *TD: n=18* **(14-20 YO)** | - | **Reduced power in L/R hemisphere (ASD vs. TD); reduced ITC in R (0.64-0.82s) and L (1.04-1.22s) hemisphere (ASD vs. TD)** | - |
| Ono, 2020 [5] | MEG: ASD vs. TD (20 & 40-Hz) | CHILDREN *ASD: n=23* *TD: n=32* **(5-7 YO)** | No difference (ASD vs. TD) | No difference (ASD vs. TD); dominance in R hemisphere (ASD, TD) | R-side 40-Hz ASSR positively correlated with age (TD only); L-side 40-Hz ASSR correlated with KABC score (ASD, TD) |
| Edgar, 2016 [6] | MEG: ASD vs. TD (40-Hz) | CHILDREN *ASD: n=42* *TD: n=48* **(7-14 YO)** | - | No difference in ASSR power or ITC (ASD vs. TD); dominance in R hemisphere (ASD, TD) | R/L-side ASSR ITC positively correlated with age by ~0.01/year (ASD, TD) |
| Rojas, 2008 [7] | MEG: ASD vs. PARENT vs. TD (40-Hz) | ADULTS *ASD: n=11* *ASD parents: n=16* *TD: n=16* **(>18 YO)** | - | **Higher induced gamma power, reduced evoked gamma power & ITC (ASD/ASD parents vs. TD)** | - |
| Wilson, 2007 [8] | MEG: ASD vs. TD (40-Hz) | CHILDREN/ ADOLESCENTS *ASD: n=10* *TD: n=10* **(7-17 YO)** | - | **Reduced power in L hemisphere (ASD vs. TD)** | - |

**Supplementary Table 1) Summary of previous studies evaluating ASSR in ASD vs. TD participants**

Significant group differences highlighted in **bold**.
Abbreviations: ASSR – auditory steady state response; ASD – autism spectrum disorder; TD – typically developing; MEG – magnetoencephalography; ITC – inter-trial coherence; KABC – Kaufman Assessment Battery for Children; L – left; R – right; YO – years old
