## Supplementary Figure 1 for "Testing the auditory steady-state response (ASSR) to 40-Hz and 27-Hz click trains in children with autism spectrum disorder and their first-degree biological relatives: A high-density electroencephalographic (EEG) study"

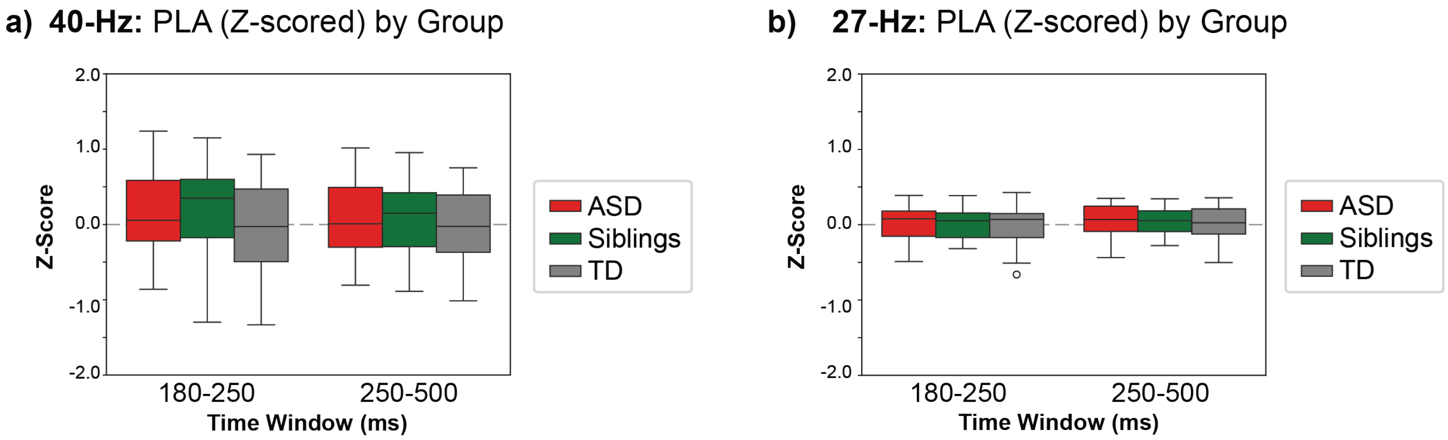


**Supplementary Figure 1) No Significant Group Differences in Phase Locking Angle**
**a-b**) phase angle difference (individual – TD mean) by z-score, comparing ASD, TD and siblings; averaged from 180-250 ms or 250-500 ms at fronto-central channels (FCz, FC3, FC4) for 40-Hz **(a)** and 27-Hz **(b)** conditions
