## Supplementary Figure 2 for "Testing the auditory steady-state response (ASSR) to 40-Hz and 27-Hz click trains in children with autism spectrum disorder and their first-degree biological relatives: A high-density electroencephalographic (EEG) study"

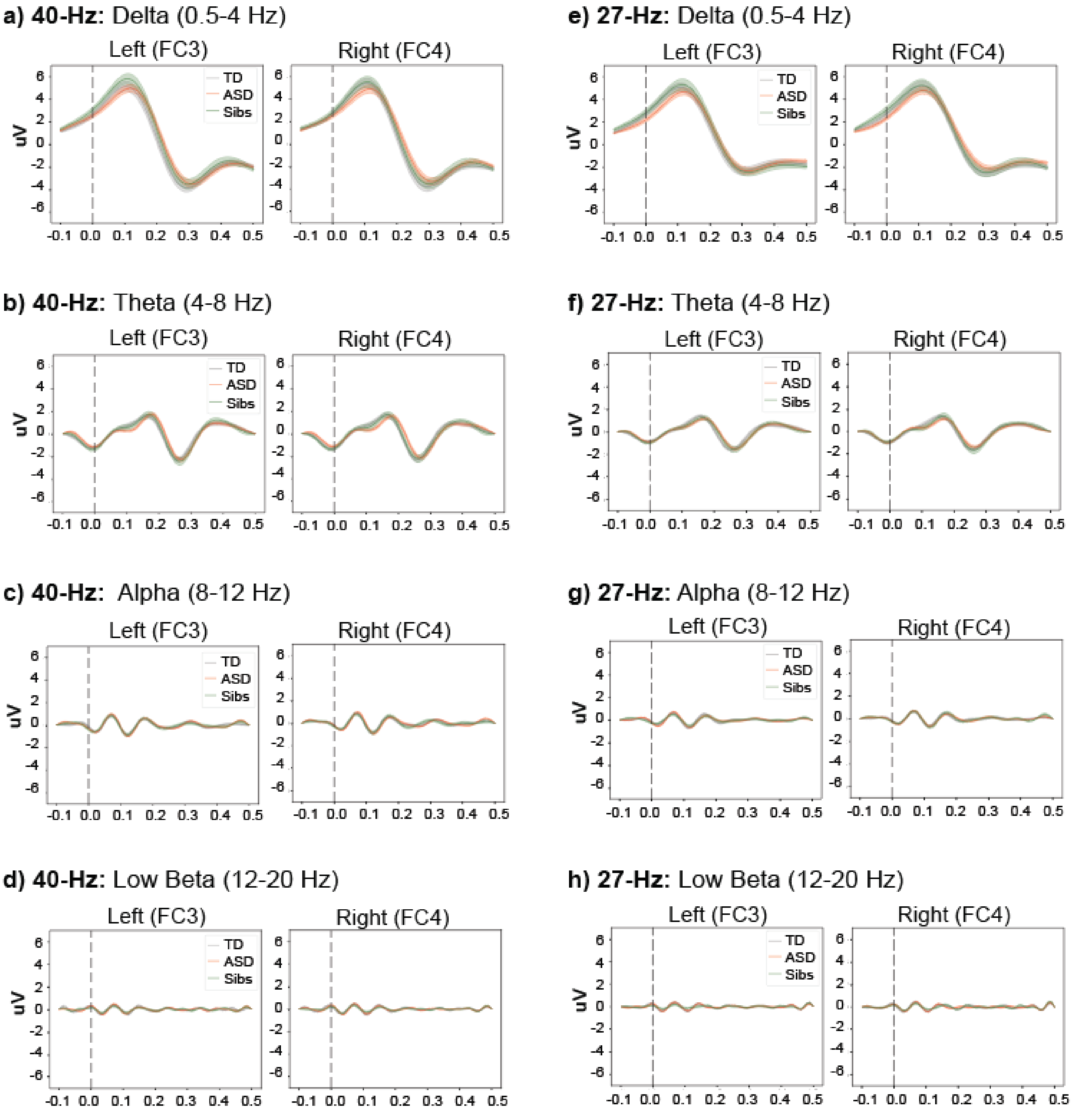


**Supplementary Figure 2) 40 and 27-Hz Evoked Response in Various Frequency Bands**
Band-pass filtered evoked response to 40-Hz **(a-d)** and 27-Hz **(e-h)** stimulation, separated by group (TD: grey, ASD: red, siblings: green) and neuro-oscillatory band (delta: 0.5-4Hz, theta: 4-8 Hz, alpha: 8-12 Hz, low beta: 12-20 Hz).
