## Supplementary Figure 3 for "Testing the auditory steady-state response (ASSR) to 40-Hz and 27-Hz click trains in children with autism spectrum disorder and their first-degree biological relatives: A high-density electroencephalographic (EEG) study"

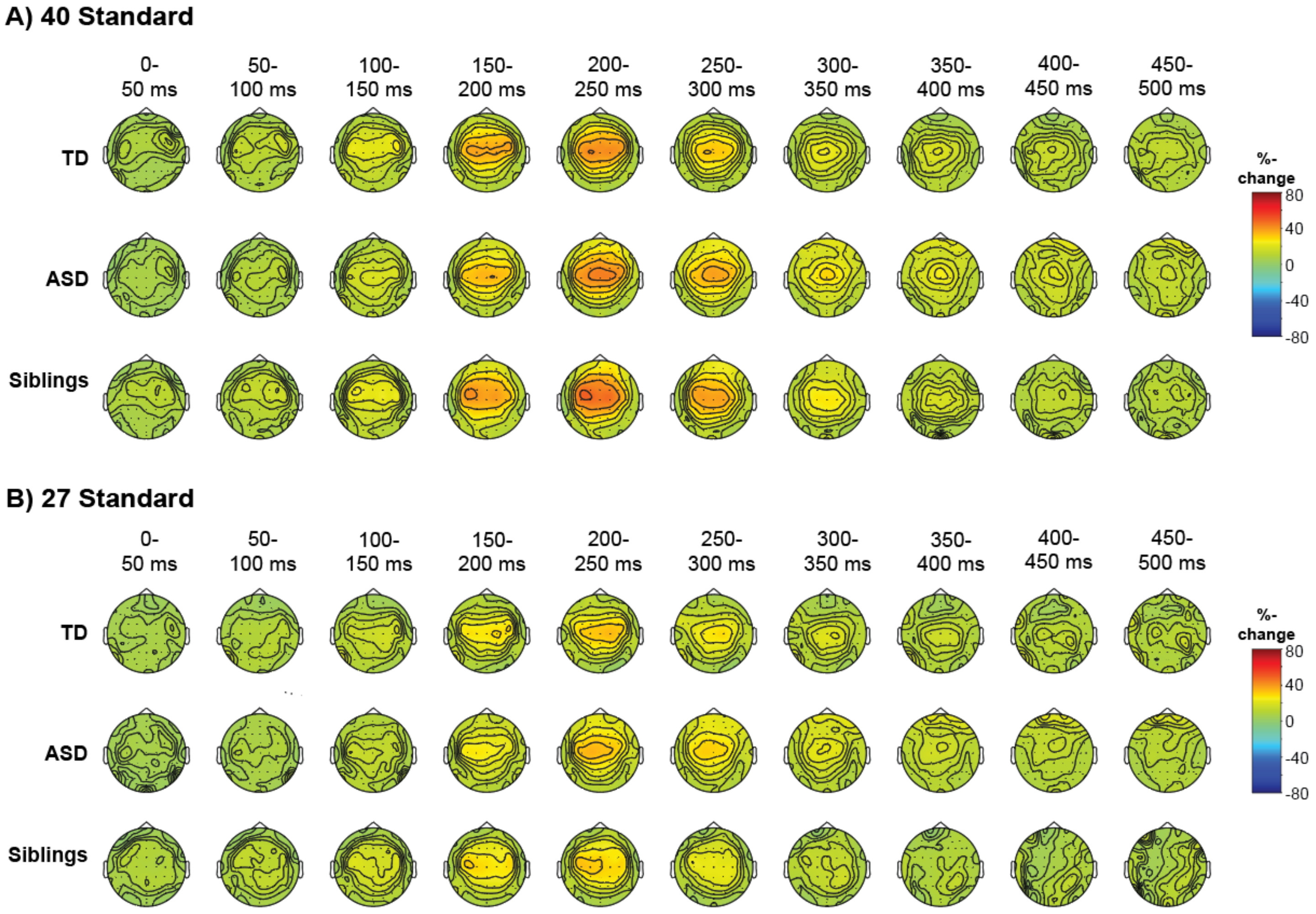


**Supplementary Figure 3) Low-Frequency [0.5-8 Hz] Topography**
Low-frequency topography (represented as %-change power relative to baseline) during 40-Hz **(a)** and 27-Hz **(b)** stimulus presentation in 50 ms windows, separated by group (TD: top, ASD: middle; siblings: bottom).
